# Glycyrrhizin inhibits LPS-induced inflammatory responses in goat ruminal epithelial cells *in vitro*

**DOI:** 10.1101/2020.05.27.118570

**Authors:** Xuanxuan Pu, Pattygouri Mullahred, Junfeng Liu, Xuefeng Guo, Jian Gao, Xiuping Zhang, Chenyu Jiang, Sujiang Zhang

**Author notes:** Xuanxuan Pu and Pattygouri Mullahred equally contributed to this work.

## Abstract

A long-term of high concentration feeding in ruminants can bring huge economic profits, but it also impose ruminants into great threat of suffering subacute ruminal acidosis (SARA). SARA is a kind of disease which attenuate the health, feed intake and production of ruminants, and when ruminants suffer SARA, the concentration of lipopolysaccharide (LPS) increase largely. Glycyrrhizin is reported to have anti-inflammation effects, and the study was conducted to investigate effects of glycyrrhizin on LPS-induced goat ruminal epithelial cells (GRECs) to provide evidence for using glycyrrhizin as a treatment for SARA. Effects of LPS, and glycyrrhizin on cell viability of GRECs were investigated, respectively. Then GRECs were stimulated with LPS (50 mg/L) for 2 h, and glycyrrhizin were added at the concentration of 0, 50, 75, 100 and 125 mg/L for 24 h to investigate the expression of inflammatory cytokines (by Elisa kits), the mRNA expression of NF-κB and inflammatory cytokines (by qRT-PCR), the distribution of Zo-1 and Occludin (by immunofluorescence staining), the expression of Occludin (by Western blot analysis), and the morphology of GRECs. The results showed that: (1) Glycyrrhizin at the concentration of 50, 75, 100, and 125 mg/L had no cytotoxic effects on GRECs, and LPS at the concentration of 50 mg/L significantly decreased the cell viability of GRECs. (2) Glycyrrhizin attenuated the expression and relative mRNA expression of TNF-α, IL-1β, IL-6, IL-8 and IL-12 by a dose-dependent manner, and significantly attenuated the relative mRNA expression of NF-κB. (3) Immunofluorescence staining and Western blot analysis showed that the quantity of Zo-1 and Occludin, and the expression of Occludin all increased with the treatment of glycyrrhizin. (4) Glycyrrhizin attenuated LPS-induced autophagy and protected the structural integrity of GRECs. In conclusion, glycyrrhizin significantly inhibited the inflammatory response in LPS-stimulated GRECs, and it may be used as a potential agent for the treatment of SARA.

## Introduction

SARA has become one of the most harmful and common diseases in ruminants breeding as more concentration were fed for the growing demand of animal productions (Hong et al., 2010). Researches showed that the incidence of ruminants’ SARA in Italy, Ireland, Netherlands and the United States were 33%, 13.8%, 11% and 19%-20.1%, respectively (Morgante et al., 2007; Kleen et al., 2009; Luke et al., 2008; Garrett et al., 1999). Yamamoto et al. (1993) reported that the rumen of fattening cattle was in a low acid environment for a long time when it suffered SARA, and finally led to the shedding of ruminal mucosa. Ruminal epithelium play a crucial role in defensing against bacteria (Wu et al., 2013), however, SARA could damage the cell viability and barrier function of ruminal epithelial cells. Even worse, systemic inflammatory response would be aroused when toxin and inflammatory substances crossed the mucosal barrier and reached to blood circulation (Penner et al., 2010). LPS is a main toxin product of *Gram negative bacteria*, and the concentration of free LPS in the rumen increases sharply when ruminants are under SARA. LPS fastened the release of inflammatory factors such as TNF-α, IL-1β and IL-6 (Yan et al., 2014). The mechanism of that is as follows: LPS leads to the activation of TLR4 signal pathway causing the release of inflammatory cytokines. Meanwhile, the NF-κB pathway, which has key regulatory functions in inflammation and innate and adaptive immunity (Rietschel et al., 1987), was also activated by LPS, and it further hastened the release of inflammatory cytokines. Histopathological change of GRECs was observed when LPS entered into rumen tissues. Autophagy, as a survival mechanism of cells in adverse environments, can remove damaged organelle and protein to protect cells (Zhu et al., 2011). Autophagy has been demonstrated to be widely involved in various disease processes, including cardiovascular disease, cancer, and neurodegenerative diseases (Lisa et al., 2012; Poon et al., 2012; Poon et al., 2012). Autophagy was also observed in GRECs when ruminants were under SARA as well. However, excessive autophagy could lead to large death of cells. Thus, we need to find agent that could prevent and treat inflammation when ruminants were under SARA.

Glycyrrhizin, isolated from licorice, has been reported to have anti-oxidative (Haraguchi et al., 1998; Vaya et al., 1997) and anti-inflammatory effects (Yamashita et al., 2017; Gonzalez-Reyes et al., 2016). Glycyrrhizin could be used to prevent some disease and to improve meat quality (Okamoto et al., 2001; Shibayama et al., 1989). Thus, we made a hypothesis that glycyrrhizin could reduce the inflammatory response in GRECs and provide treatment for SARA. But there was less research investigating on anti-inflammatory effects of glycyrrhizin on LPS-induced GRECs. Therefore, in this study, GRECs were stimulated with LPS, and the effects of glycyrrhizin in LPS-stimulated GRECs were investigated *in vitro* to provide evidence for using glycyrrhizin as a method to treat ruminants under SARA.

## Materials and Methods

### Reagents

Glycyrrhizin (purity >99%) was purchased from Yuanye Biotechnology Co., Ltd. (Shanghai, China). LPS (Escherichia coli 055:B5) was purchased from Sigma Chemical (St. Louis, MO, USA). Fetal bovine serum were purchased from Gibco (Grand Island, New York, USA). DMEM/F12 was purchased from Hyclone (Beijing, China). Anti-cytokeratin-18 IgG and FITC-goat anti mouse IgG were purchased from Santa cruz biotechnology Co., Ltd. (America). Anti-rabbit IgG H&L was purchased from Abcam (America). DAPI were purchased from Tongren institute of chemistry (Japan). Elisa kits for TNF-α, IL-1β, IL-6, IL-8 and IL-12 were gotten from Biolegend (Camino Santa Fe, CA, USA). BCA protein quantitative kit and SYBR Green PCR kit were purchased from Smer Fell Science and Technology Co., Ltd. (Shanghai, China). RIPA, Acrylamide(29:1), Tris-HCl pH=8.8 electrophoretic buffer, Tris-HCl pH=6.8 electrophoretic buffer, SDS, TEMED, PBS phosphate buffer and DEPC were purchased from JRDUN Biotechnology Co., Ltd. (Shanghai, China). β-actin was purchased from Tianjin Sungene Biotech Co. (Tianjin, China). Alexa Fluor 488 Labeled goat anti-rabbit IgG (H+L), DAPI, Goat anti-rabbit-HRP secondary sera, Donkey anti-goat-HRP secondary sera and Goat anti-mouse-HRP secondary sera were purchased from Biyuntian biotechnology co., Ltd. (Shanghai, China). Occludin (rabbit) and ZO-1 (rabbit) were purchased from Aobasen biotechnology co., Ltd (Beijing, China). DMEM/F12 was purchased from Hyclone (Los Angeles, USA), MiniBEST Universal RNA Extraction Kit was purchased from TaKaRA, iScript™ gDNA Clear Synthesis Kit was purchased from Bio-Rad.

### Animals and treatment

All experimental procedures were approved by the Animal Research Ethics Committee of Tarim University (TLMUMO-2018-10) and all experiments followed the guidelines for the care and use of laboratory animals published by the US National Institutes (2010). Goats weighed 10 kg were fed with standard laboratory diets with free access to water in the experimental station of Tarim University.

### Cell culture, treatment and identification

Large-scale of rumen epithelial tissues of abdominal sac were collected and mixed together from goats. Primary rumianl epithelial cells were obtained as described in previous study (Zhao et al., 2019). Primary cells were purified, and were identified by the method of cytokeratin-18 immunocytochemistry.

### Cell viability assay

The cell viability of LPS, and glycyrrhizin in GRECs were measured by [3-(4,5-dimethylthiazol-2-yl)-2,5-diphenyl-tetra-zoliumbromide] (MTT) assay, respectively. The GRECs were treated with 100 μL of LPS (50 mg/L), and glycyrrhizin (0, 50, 75, 100 and 125 mg/L) for 24 h, respectively, and then 10 µL of 5 mg/L MTT was added for 4 h. Finally, the supernatants were removed and dissolved with 150 ml of DMSO in each well. Absorbance was determined at 570 nm.

### Inflammatory cytokines assay

In the following assay, GRECs were stimulated with 50 mg/L of LPS for 2 h firstly, and then glycyrrhizin (0, 50, 75, 100 and 125 mg/L) were added for 24 h. Cells were centrifuged at 750 g for 20 min and the supernate was collected. The levels of TNF-α, IL-1β, IL-6, IL-8 and IL-12 were measured using Elisa kits according to the manufacturer’s instruction.

### The relative mRNA expression of NF-κB, TNF-α, IL-1β, IL-6, IL-8 and IL-12 Primers design and synthesis

According to the mRNA sequences of NF-κB, TNF-α, IL-1β, IL-6, IL-8, IL-12 and GAPDH of goats in GenBank. The primers of mRNA were designed by primer 5.0 (Wang et al., 2015; Wang et al., 2015). Primers were synthesized by JRDUN Biotechnology Co., Ltd., (Shanghai, China) as shown in Table 1.

**Table 1.**
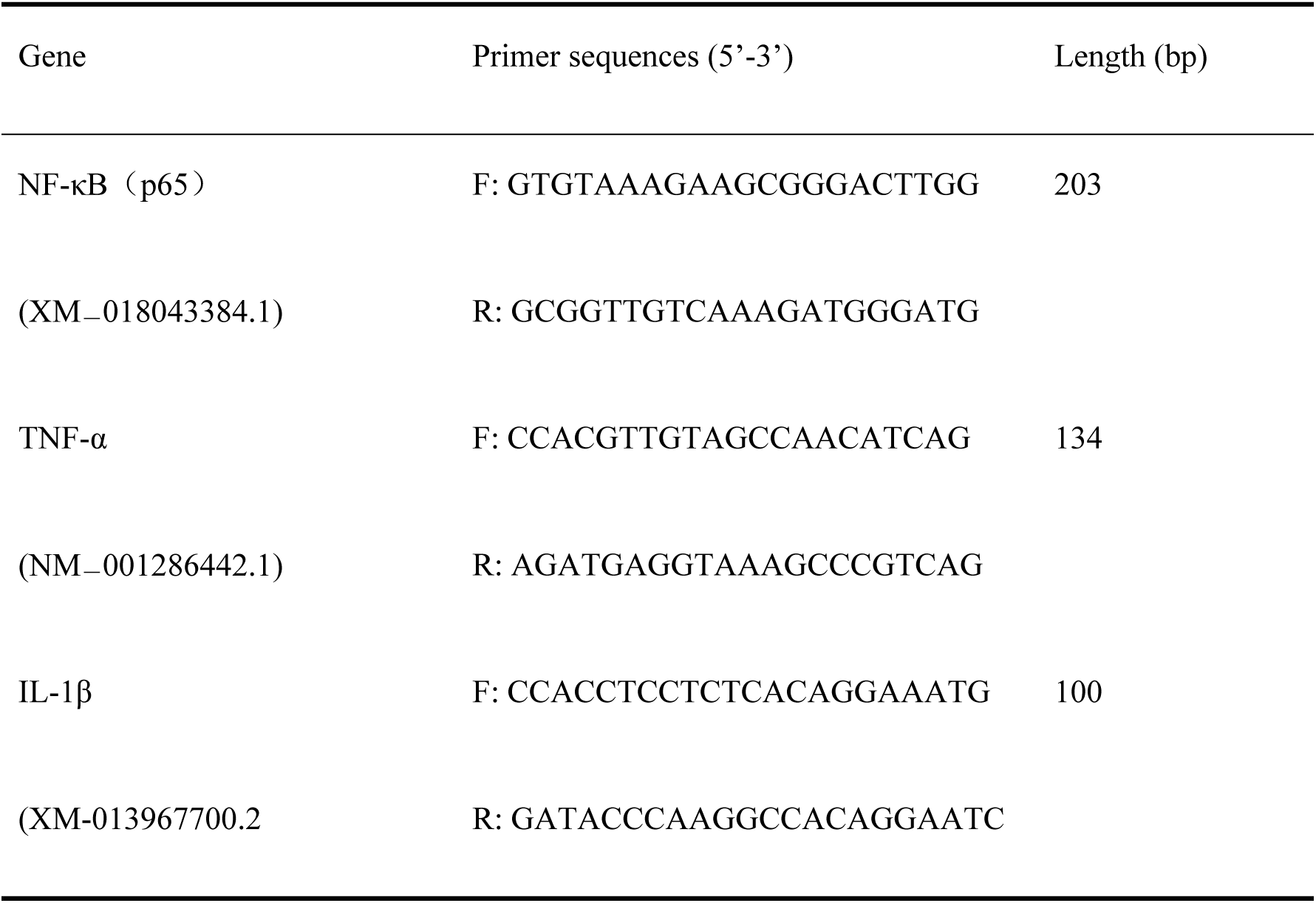
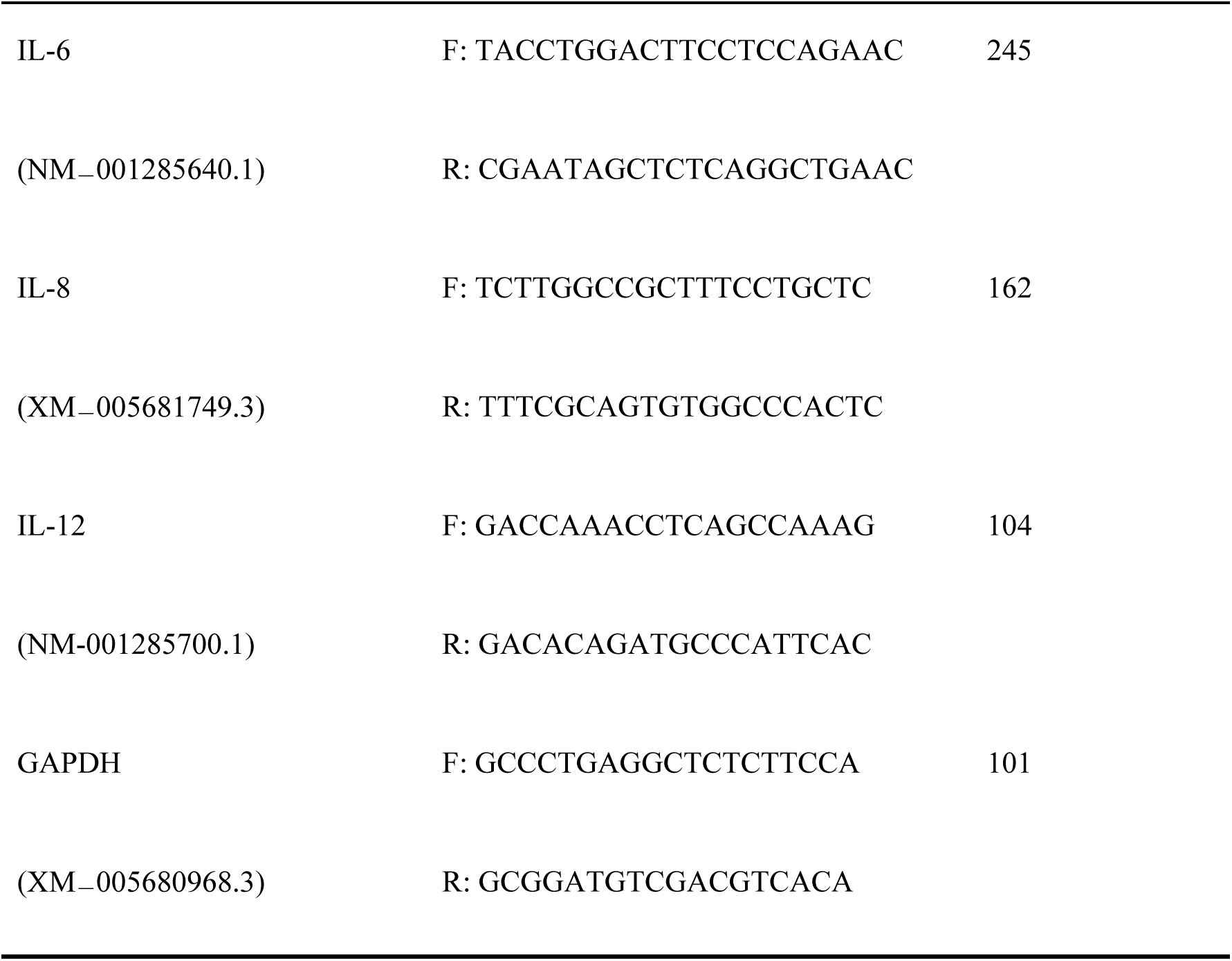
The primer sequence of targeted gene.

### RNA extraction and reverse transcription

GRECs were treated as mentioned above, the total RNA of cells was extracted by MiniBEST Universal RNA Extraction Kit according to the manufacturer’s instructions, and the production was measured by 1% agarose gel. Reverse transcription was as follows: first, DNA contamination of RNA samples was removed with a RNase-free DNase. Then cDNA was synthesized with reference to the reverse transcription kit instructions of Dalian Biotechnology Co., Ltd.

### qRT-PCR

The reversed cDNA was diluted to 10 μM and was subjected to qRT-PCR by SYBR Green PCR kit according to the manufacturer’s instruction. PCR was carried out in triplicate 25-μL reactions containing 12.5 μL SYBRGreen Mix, 0.5 μL upstream primer F, 0.5 μL downstream primer R, 9.5 μL ddH_2_O, 2 μL cDNA template. The response procedures were: 95 °C, 10 min (95 °C, 15 s; 60 °C, 45 s) × 40; 95 °C, 15 s; 60 °C, 1 min; 95 °C, 15 s; 60 °C, 15 s.

### The distribution and quantity of Occludin and Zo-1

The distribution and quantity of Occludin and Zo-1 was measured by immunofluorescence staining. GRECs were treated as mentioned above, the slides of GRECs were washed with 0.02 M PBS in the culture plate, fixed by 4% paraformaldehyde for 30 min, permeabilized by 0.1% Triton X-100, blocked by 1% bovine serum albumin (BSA) for 1 h, and each step followed three times washing by 0.02 M PBS. Cell slides was incubated overnight at 4°C in a wet box with a specific primary antibody (1:50), and cell slides were washed by 0.02 M PBS for three times. Then cell slides were incubated in a wet box at room temperature for 1 h with the added secondary antibody labeled with fluorescein isothiocyanate (1:500), and then washed by 0.02 M PBS for three times, and afterwards were sealed by adding anti-quenching seal tablets and DAPI at a 1:500 ratio, stored in a -20 °C, and photographed by a fluorescence microscope.

### Western blot analysis

Approximately 5×10^5^ GRECs (treated as mentioned above) of each group were added into Lysate with protease and phosphatase inhibitors at 4 °C to disrupt cells thoroughly, cuffed the cells into a 1.5 mL EP tube and heated them at 95 °C for 10 min, centrifuged at 12000 g for 10 min, then the supernate was collected and the protein was quantified by the standard curve (the standard curve was measured In our study). The proteins were separated by SDS/PAGE and electrophoretically transferred onto a semi-dry rotating membrane. The membrane was blocked in NaCl/Tris containing 5% nonfat dry milk at room temperature for 1 h, and incubated with a primary antibody at 4 °C for 12 h and then incubated with a secondary antibody at 37 °C for 1 h. Finally, the blots were developed with the ECL Plus Western Blotting Detection System (GE Healthcare, Chalfont St Giles, UK).

### Morphology change of LPS-induced GRECs

To further identify anti-inflammatory effects of glycyrrhizin, the morphology of GRECs was investigated as well. GRECs treated as mentioned above were washed by PBS, resuspended and sealed by 2.5% glutaraldehyde for 2 h and washed by PBS for three times. Then tissues were sealed by osmic acid for 2.5 h and washed by PBS for three times. Tissues were dehydrated with different concentration ethanol of 30%, 50%, 70%, 90%, 95% and 100%, each dehydration lasted for 10-15 min for three times. Then tissues were replaced with propylene epoxide for two times, each time last for 10 min. Tissues were permeated by propylene oxide and epon resinand at a 2:1 ratio and by propylene oxide and epon resinand at a 1:2 ratio for 2 h, respectively, then tissues were permeated by whole epon resinand for the night. Tissues were polymerizated at 37 °C for 12 h, 45 °C for 24 h and 60 °C for 48 h, and finally, the tissues were made into cell slice and observed by electron microscopic.

### Statistical analysis

All data was presented as the means ± S.E.M by SPSS 16.0. Difference were performed by one-way ANOVA combined with Turkey’s multiple comparison tests. Differences were considered to be significant at *P*< 0.05 (*), extremely significant at *P*< 0.01 (**).

## Results

### The cytokeratin-18 expression of cells

The purified epithelial cells were identified by the detection of cytokeratin-18, and the result was shown in Fig. 1. Green fluorescence was showed in our result, and the positive staining for cytokeratin-18 indicated that the purified ruminal epithelial cells were GRECs.

**Fig. 1.**
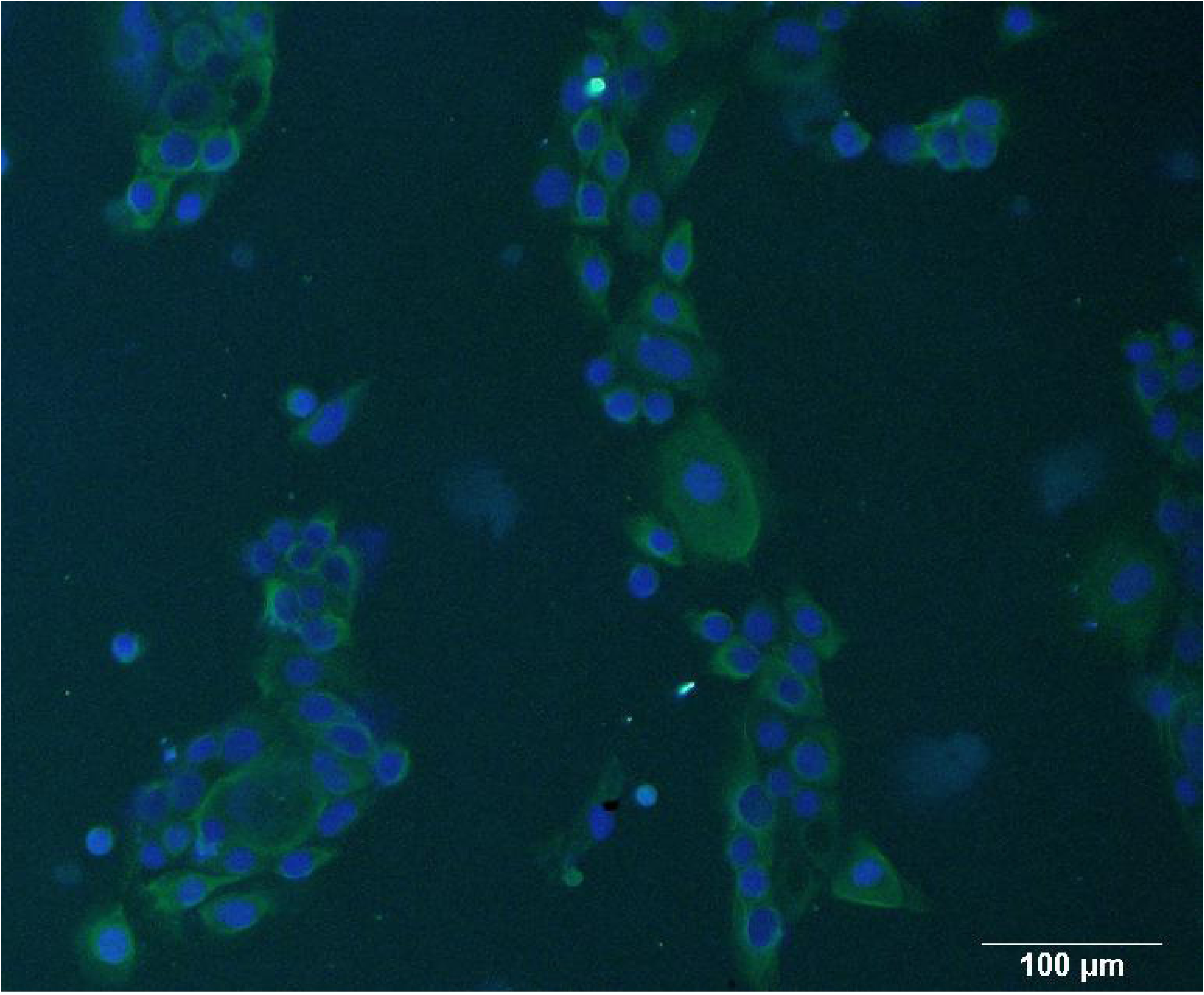
Immunocytochemistry of purified epithelial cells. Purified epithelial cells were obtained by digestion, and identified by the detection of cytoskeleton 18.

### Effects of LPS and glycyrrhizin in cell viability of GRECs

As shown in Fig. 2, the cell viability in groups added with glycyrrhizin at the concentration of 50, 75, 100 and 125 mg/L had no significant difference versus control group. The cell viability of GRECs stimulated by LPS was significantly decreased.

**Fig. 2.**
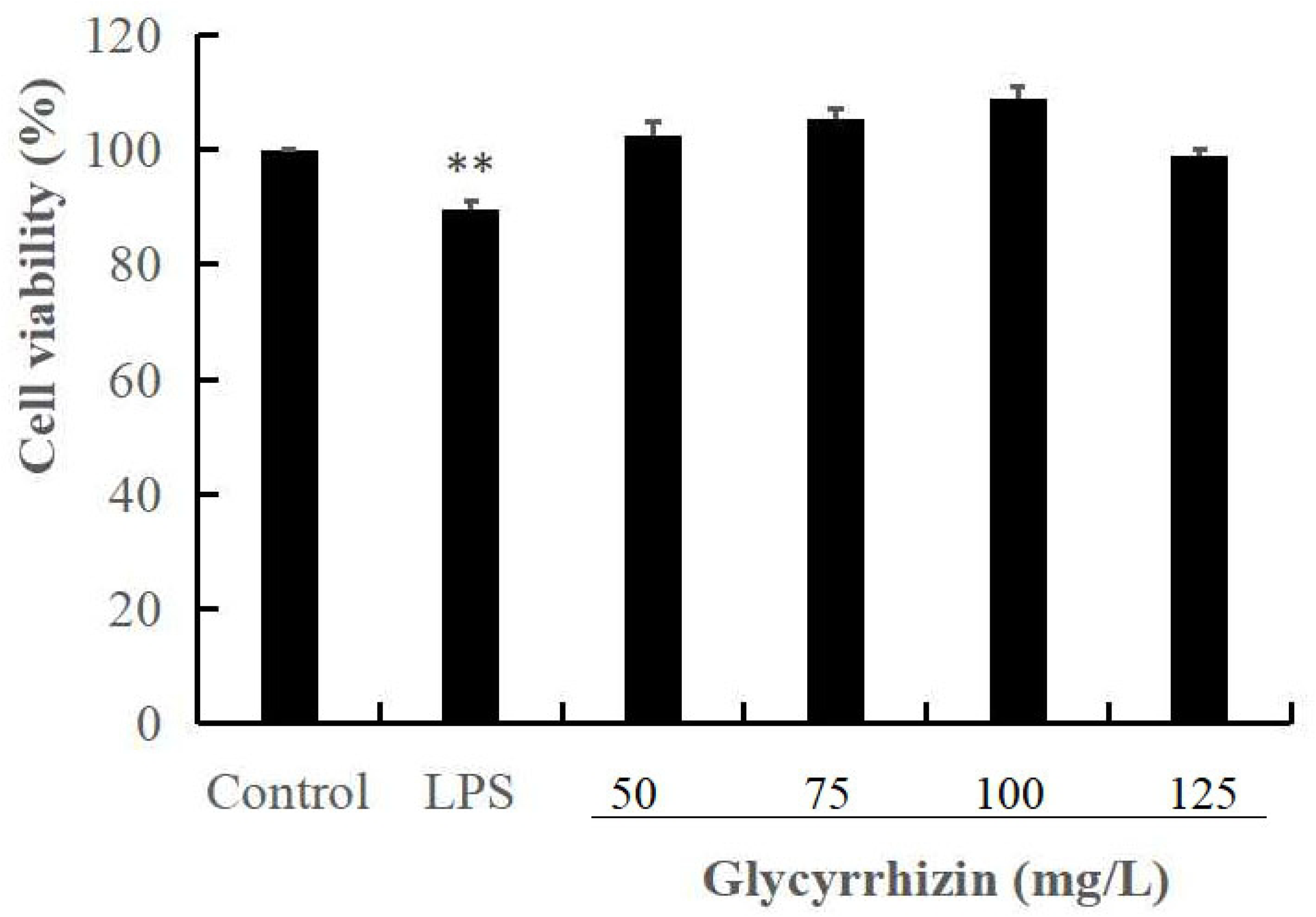
Effects of LPS and glycyrrhizin on cell viability of GRECs. The cell viability was measured by MTT assay. GRECs were incubated with LPS at the concentration of 50 mg/L, and glycyrrhizin at the concentration of 0, 50, 75, 100 and 125 mg/L, respectively. The data presented was the means ± S.E.M of three independent experiments.** *P*<0.01, * *P*<0.05 versus the control group.

### Effects of glycyrrhizin on inflammatory cytokines in LPS-induced GRECs

As shown in Fig. 3, the production of TNF-α, IL-1β, IL-6, IL-8 and IL-12 in LPS group were extremely significant higher than those with the treatment of glycyrrhizin (*P*<0.01). Glycyrrhizin at the concentration of 50, 75, 100 and 125 mg/L inhibited the production of cytokines by a dose-dependent manner.

**Fig. 3.**
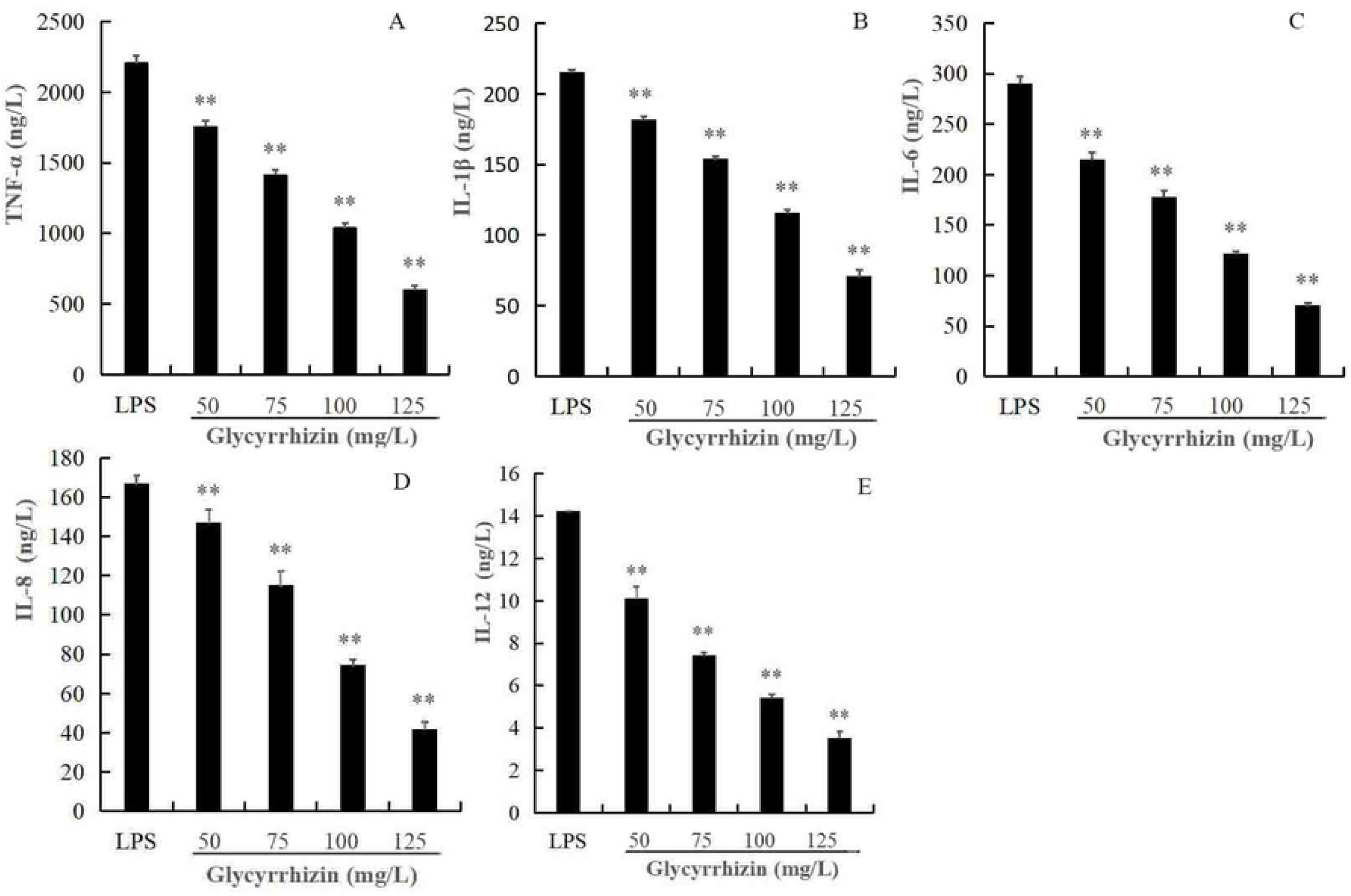
**(A-E)**. Effects of glycyrrhizin on LPS-induced inflammatory cytokines. GRECs were incubated with 50 mg/L of LPS for 2 h, and then glycyrrhizin (0, 50, 75, 100, 125 mg/L) was added for 24 h. Inflammatory cytokines of TNF-α (A), IL-1β (B), IL-6 (C), IL-8 (D) and IL-12 (E) were measured by Elisa kits. The data presented were the means ± S.E.M of three independent experiments. ** *P*<0.01, * *P*<0.05versus LPS group.

### Effects of glycyrrhizin on the relative mRNA expression of NF-κb, TNF-α, IL-1β, IL-6, IL-8 and IL-12 in LPS-induced GRECs

As shown in Fig. 4, glycyrrhizin at the concentration of 50, 75, 100 and 125 mg/L inhibited the mRNA expression of TNF-α, IL-1 β, IL-6, IL-8 and IL-12 in LPS group by a dose-dependent manner (*P*<0.01). Glycyrrhizin added at the concentration of 75, 100 and 125 mg/L decreased the mRNA expression of NF-κb significantly.

**Fig. 4.**
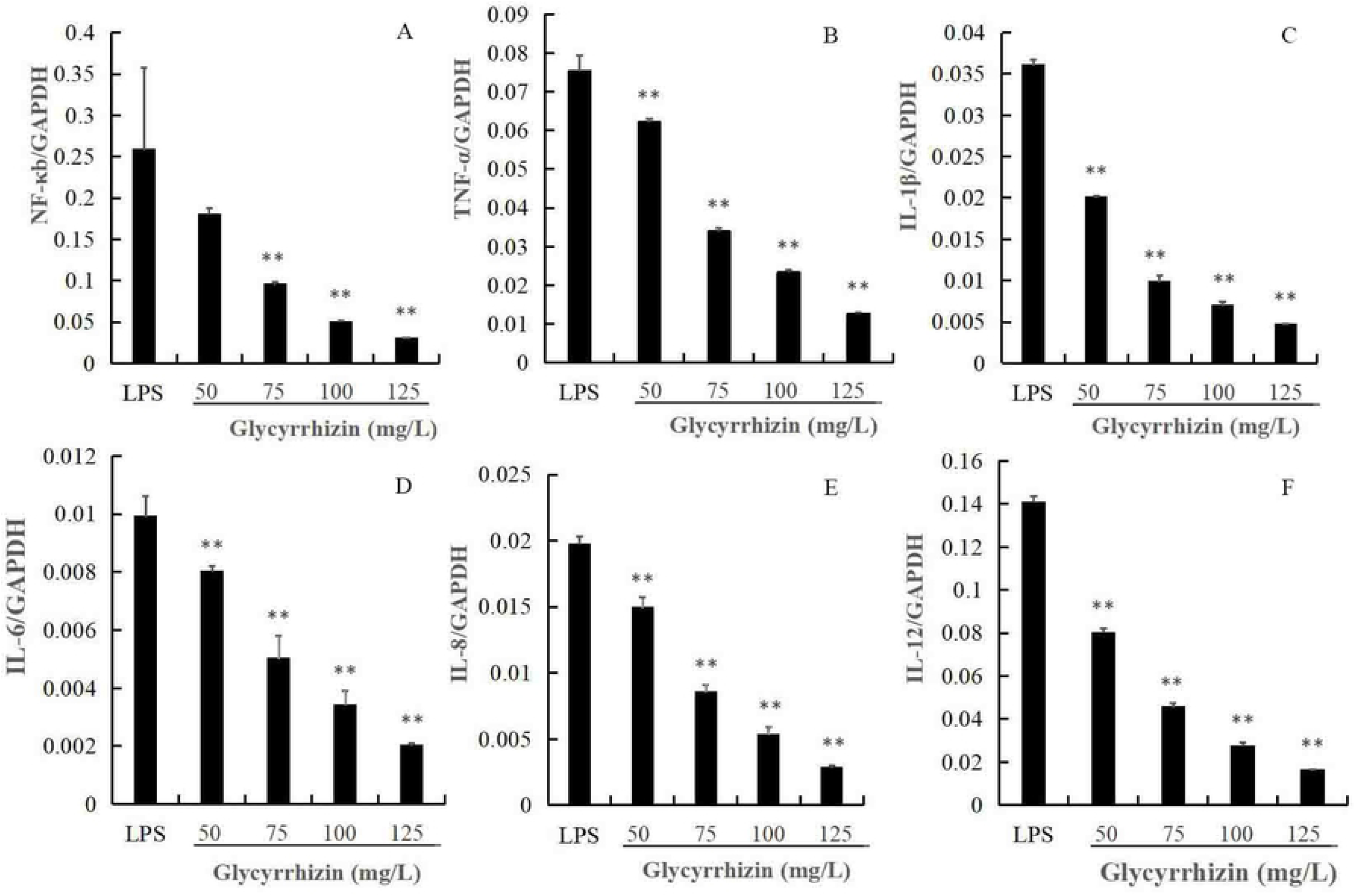
**(A-F)**. Effects of glycyrrhizin on the relative mRNA expression of NF-κB, TNF-α, IL-1 β, IL-6, IL-8 and IL-12 in LPS-induced GRECs. GRECs were stimulated with 50 mg/L of LPS for 2 h, and then were incubated with glycyrrhizin (0, 50, 75, 100, 125 mg/L) for 24 h. The relative mRNA expression of NF-κb (A), TNF-α (B), IL-1β (C), IL-6 (D), IL-8 (E) and IL-12 (F) was measured by qRT-PCR. The data presented were as the means ± S.E.M of three independent experiments. ** *P*<0.01, * *P*<0.05 versus LPS group.

### Effects of glycyrrhizin on the distribution of Zo-1 and Occludin in LPS-induced GRECs

The distribution of Occludin and Zo-1 in GRECs was detected by immunofluorescence staining. As shown in Fig. 5, glycyrrhizin increased the quantity of Occludin and Zo-1 versus control group, and their distribution became more uniform with higher concentration of glycyrrhizin.

**Fig. 5.**
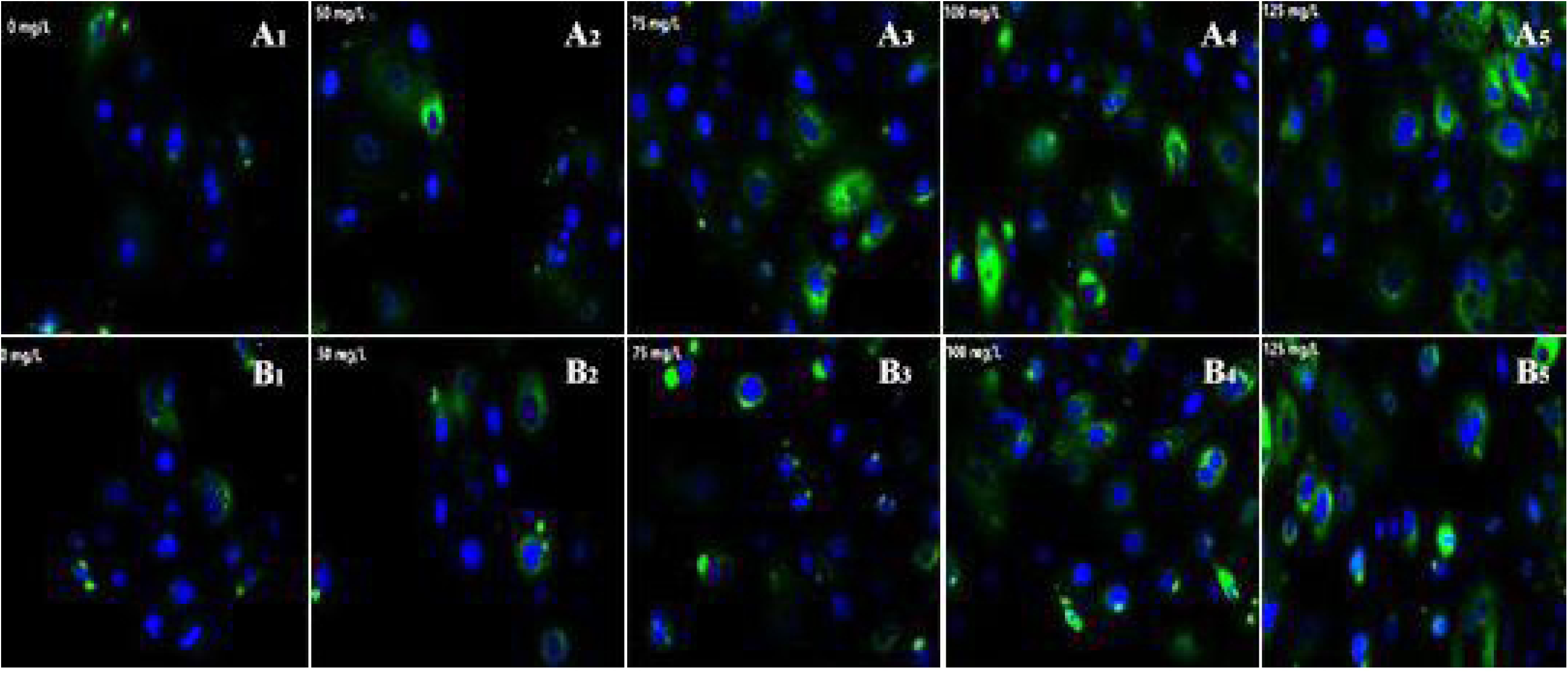
**(A-B)**. Effects of glycyrrhizin on the distribution of Zo-1 and Occludin in LPS-induced GRECs. GRECs were stimulated with 50 mg/L of LPS for 2 h, and then were incubated with glycyrrhizin (0, 50, 75, 100, 125 mg/L) for 24 h. The distribution of the Zo-1 (A_1_-A_5_) and Occludin (B_1_-B_5_) was measured by immunofluorescence staining. Fig. 5 (A_1_-A_5_) and Fig. 5 (B_1_-B_5_) represented the LPS group and LPS-induced GRECs treated with glycyrrhizin at the dose of 50, 75, 100, 125 mg/L, respectively.

### Effects of glycyrrhizin on the expression of Occludin in LPS-induced GRECs

As shown in Fig. 6(A-B), the expression of Occludin in LPS group was extremely significant lower than those with the treatment of glycyrrhizin (*P*<0.01), and the expression of Occludin increased with the treatment of glycyrrhizin by a dose-dependent manner.

**Fig. 6.**
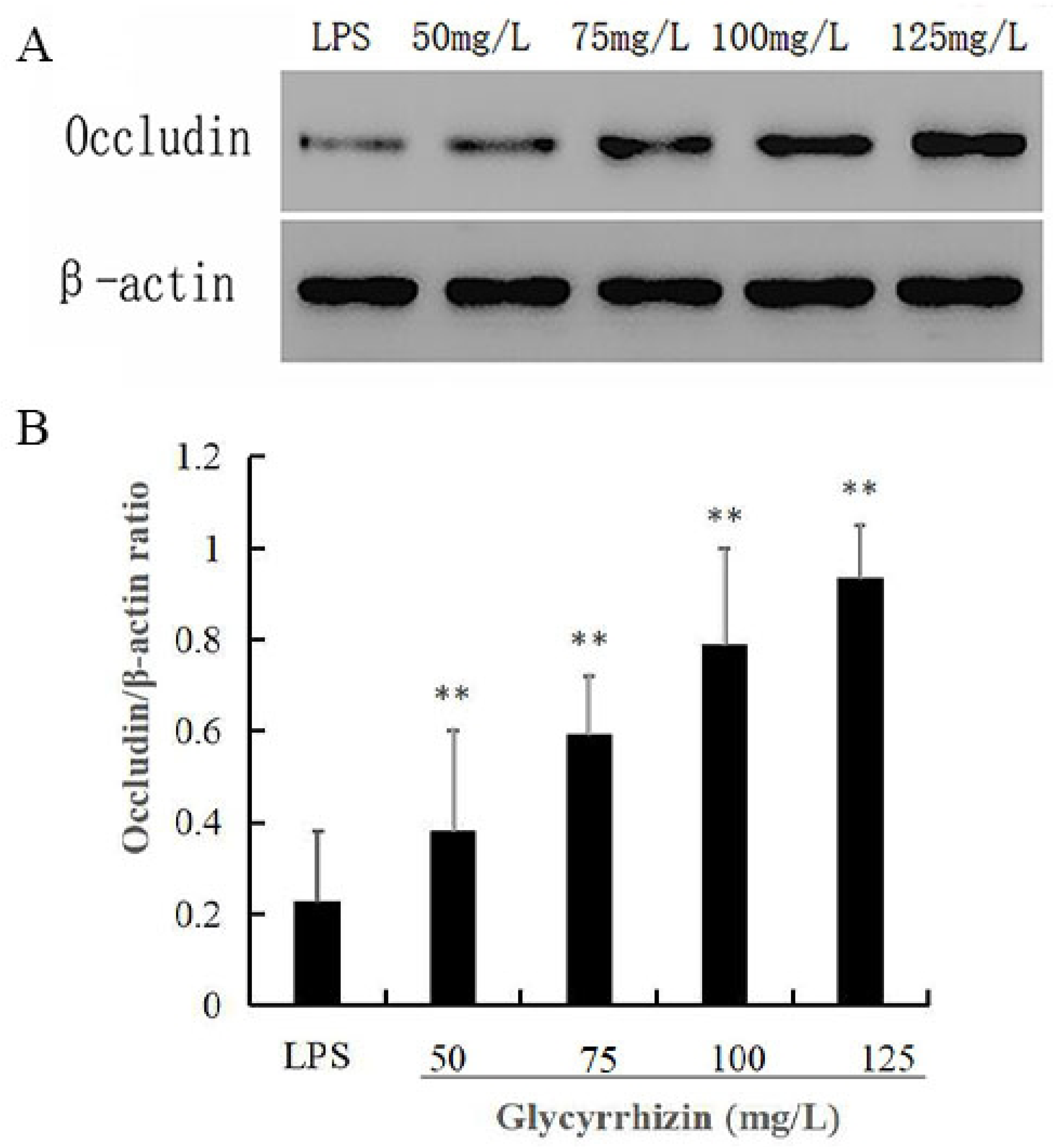
**(A-B)**. Effects of glycyrrhizin on the expression of Occludin in LPS-induced GRECs. GRECs were incubated with 50 mg/L of LPS for 2 h, and then were incubated with glycyrrhizin at 0, 50, 75, 100, 125 mg/L for 24 h. The expression of Occludin was measured by Western blot analysis. β-actin was used as a control, the values presented were the means ± S.E.M of three independent experiments. ** *P*<0.01, * *P*<0.05 versus LPS group.

### Effect of glycyrrhizin on the morphology of LPS-induced GRECs

As shown in Fig. 7A, LPS group was with high electron density in cytoplasm, the autophagy around the nucleus were the highest and most of it was with high electron density. As shown in Fig. 7C, Fig. 7D and Fig. 7E, the pathological changes were reduced with added glycyrrhizin. The autophagy around the nucleus occured significantly in LPS group, while its quantity decreased significantly in Fig. 7C, Fig. 7D and Fig. 7E. What’s more, the autophagy with high electron density was decreased with glycyrrhizin by a dose-dependent manner as shown in Fig. 7C, Fig. 7D and Fig. 7E. Thus better cellular morphology of GRECs was founded with higher treatment of glycyrrhizin.

**Fig. 7.**
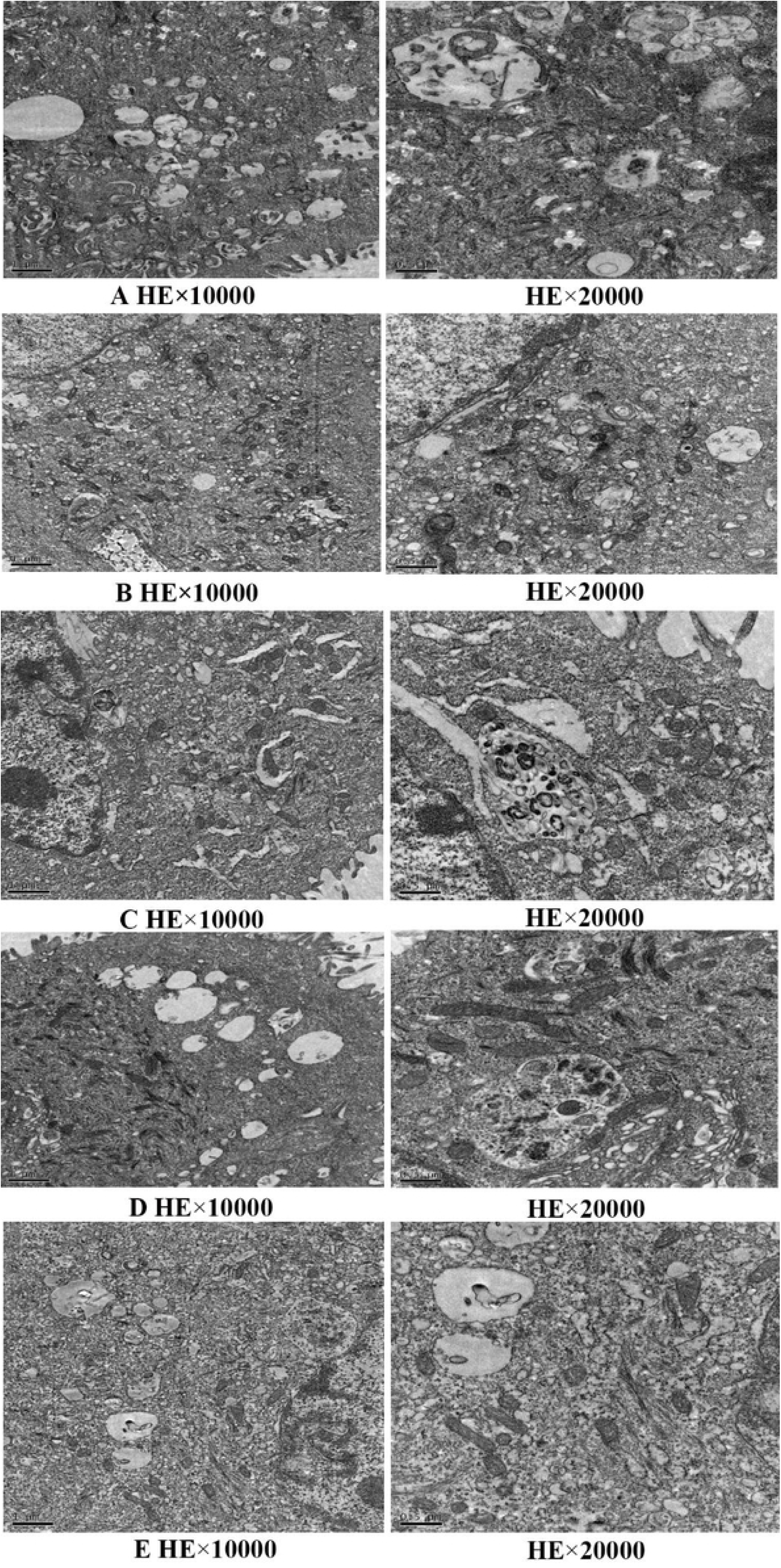
**(A-E)**. Effects of glycyrrhizin on morphology of LPS-induced GRECs. GRECs were incubated with LPS for 2 h, and glycyrrhizin was added at the dose of 0, 50, 75, 100, 125 mg/L from picture A to E. The autophagy around the nucleus was the highest in fig. A, and it gradually decreased in fig. C, fig. D and fig. E.

## Discussion

### Cell viability of glycyrrhizin in GRECs

When ruminants were under SARA, the rumen mucosal barrier was impaired, and the content of LPS increased sharply (Rodríguez-Lecompte et al., 2014; Khafipour et al., 2009). The mRNA expression of inflammatory cytokines was increased significantly in LPS-induced ruminal epithelium of dairy cattle (Zhang et al., 2016). The tight connection between GRECs was destroyed (Liu et al., 2013), and the inflammatory cytokines would enter into the rumen leading to rumen diseases such as rumen acidosis and the death of the ruminants. Glycyrrhizin which has anti-inflammatory effects in many disease may be used as a potential anti-inflammatory agent for the treatment of SARA. Wang et al (2017) showed that glycyrrhizin at the concentration of 50, 100, and 200 μg/ml had no cytotoxic effects on mouse endometrial epithelial cells. In our assay, glycyrrhizin (0-125 mg/L) had no cytotoxic effects on GRECs as well.

### Effects of glycyrrhizin in LPS-induced GRECs

It has been reported that glycyrrhiza glabrathe had anti-inflammatory and antiviral effects (Uto et al., 2012). Glycyrrhizin has been reported to inhibit LPS-induced inflammatory cytokines in many cells (Wang et al., 2017; Li et al 2010). Fei et al., (2014) showed that glycyrrhizin decreased the content of TNF-α and IL-6 significantly in caerulein-induced acute pancreatitis of mice. Kong et al., (2019) showed that glycyrrhizin decreased the content of IL-4 and IL-6 a lot in LPS-induced acute lung injury of mice. Fu et al., (2014) reported glycyrrhizin inhibited the expression of factor-α and IL-6 *in vitro*. Similarly, in our study, glycyrrhizin dose-dependently inhibited the production of TNF-α, IL-1β, IL-6, IL-8 and IL-12.

NF-κB is considered to be an important pathogenic factor in many acute and chronic inflammatory diseases (Wullaert et al., 2011), and the activation of NF-κB is a central link in inflammatory reaction. NF-κB signaling pathway could regulate transcription of infammatory cytokines (Oeckinghaus et al., 2009). When NF-κB was activated by LPS, the expression of inflammatory cytokines increased (Bruewer et al., 2003; Wellnitz et al., 2011), and these inflammatory cytokines could further activate NF-κB. Stimulating ruminal epithelial cells with LPS could lead to the activation of NF-κB signaling pathway (Schaefer et al., 2004), and in our study, LPS at the concentration of 50 mg/L had significantly decreased the cell viability of GRECs. but Strandberg et al., (2005) showed that LPS of 50 mg/L had no significant effect on bovine primary mammary epithelial cells. Wellnitz et al., (2004) reported that 200 mg/L LPS could affect the primary mammary epithelial cells of Holstein cows. This difference might be caused by the difference of animals and method used. In our study, the results showed that glycyrrhizin dose-dependently inhibited the relative mRNA expression of NF-κB, TNF-α, IL-1β, IL-6, IL-8 and IL-12, which indicate that the inhibition of NF-κB signaling pathway might lead to the decreased production of inflammatory cytokines, and it needs further investigation. TNF-α is an anti-tumor multi-function factor and large amounts of TNF-α can cause serious inflammatory reactions like tissue necrosis. Research showed that TNF-α could inhibit the expression of Occludin largely (Song et al., 2004; Bruewer et al., 2003; Xu et al., 2010). Therefore, if the expression of Occludin was increased in LPS-induced GRECs, it could further identify the anti-inflammatory effects of glycyrrhizin.

To further identify anti-inflammatory effects of glycyrrhizin, the distribution of tight junction proteins and the expression of Occludin were investigated in our study. The rumen epithelium from mucous layer to serosa layer was stratum corneum, stratum granulosum, stratum spinosum and stratum basale which play an important role in protection. There are tight connection proteins in stratum granulosum and they play an important role in maintaining rumen barrier function (Harhaj et al., 2004; Steele et al., 2011). The tight junction proteins are composed of Claudin, Occludin, Zo-1, ZOs and so on. Sun et al., (2018) reported that when the epithelial cells were under SARA, rumen epithelium and the mRNA expression of tight junction were affected by LPS *in vitro*. When the ruminal epithelial cells were induced by LPS, the tight junction proteins were destroyed, and the cell gap became larger. In our study, the distribution of Occludin and Zo-1 became uniform and their quantity increased with the treatment of glycyrrhizin. What’s more, glycyrrhizin dose-dependently increased the expression of Occludin. The results further identified anti-inflammatory effects of glycyrrhizin in LPS-induced GRECs.

In our study, there was autophagy occurred in LPS-induced GRECs. Autophagy can be induced by many factors (Chen et al., 2008; Wang et al., 2010). Xu et al., (2006) reported that LPS could induce macrophages to produce autophagy. Abnormal activation of autophagy is closely associated with the development of acute pancreatitis (Kang et al., ;Zhang et al., 2011). In our study, glycyrrhizin attenuated the LPS-induced autophagy and dose-dependently decreased the autophagy with high electron density. Therefore, glycyrrhizin inhibited the abnormal activation of autophagy in LPS-induced GRECs.

Taken together, glycyrrhizin dose-dependently inhibited LPS-induced inflammatory response in our study. Therefore, higher concentration of glycyrrhizin should be investigated for further study to identify the upper limit concentration of glycyrrhizin.

## Conclusion

Glycyrrhizin (0-125 mg/L) had no cytotoxic effects on GRECs, and LPS significantly decreased the cell viability of GRECs. Glycyrrhizin inhibited the inflammation response in LPS stimulated GRECs by a dose-dependent manner by inhibiting the expression of inflammatory cytokines, suppressing NF-κB signaling pathway, increasing the expression of Occludin and reducing autophagy in LPS-induced GRECs. The comprehensive results showed that glycyrrhizin might be a valuable agent for the treatment of SARA.

## Author contributions

Contributed reagents/materials/analysis/ tools: X.G.; Performed the experiments: X.P; Analyzed the data: X.P; Writing-original draft: X.P.; Writing–review editing: X.P., X.G., C.J., J.L., X.Z., and S.Z..

## Acknowledgments

We acknowledge Dr Long Cheng from the University of Melbourne for useful discussion. This work was supported by the Project of Mr. Yang Sheng’s Student Community Research (2016A20006) and Xinjiang Production and Construction Group with the young and middle-aged innovation talents fund (2016BC001).

